# A simple cloning-free method to efficiently induce gene expression using CRISPR/Cas9

**DOI:** 10.1101/400838

**Authors:** Lyujie Fang, Sandy S.C. Hung, Jennifer Yek, Tu Nguyen, Shahnaz Khan, Alex W Hewitt, Raymond C.B. Wong

## Abstract

Gain-of-function studies often require the tedious cloning of transgene cDNA into vectors for overexpression beyond the physiological expression levels. The rapid development of CRISPR/Cas technology presents promising opportunities to address these issues. Here we report a simple, cloning-free method to induce gene expression at endogenous locus using CRISPR/Cas9 activators. Our strategy utilises synthesized sgRNA expression cassettes to direct a nuclease-null Cas9 complex fused with transcriptional activators (VP64, p65 and Rta) for site-specific induction of endogenous genes. This strategy allows rapid initiation of gain-of-function studies in the same day. Using this cloning-free approach, we tested two CRISPR activation systems, dSpCas9VPR and dSaCas9VPR, for induction of multiple genes in human and rat cells. Our results showed that both CRISPR activators allow efficient induction of six different neural development genes (*CRX, RORB, RAX, OTX2, ASCL1* and *NEUROD1*) in human cells, whereas the rat cells exhibit a more variable and less efficient levels of gene induction, as observed in three different genes (*Ascl1, Neurod1, Nrl*). Altogether, this study provides a simple method to efficiently activate endogenous gene expression using CRISPR/Cas9 activators, which can be applies as a rapid workflow to initiate gain-of-function studies for a range of molecular and cell biology disciplines.

## Introduction

Conventional gene overexpression studies require the need to clone the transgene cDNA into an expression vector, and therefore involves DNA ligation, bacterial transformation, screening clones, plasmid purification and quality check to confirm the vector sequences. This represents a tedious and costly procedure, especially for large-scale genome-wide overexpression study. Further to this, the use of whole transgene cDNA imposes a challenge for overexpressing multiple genes simultaneously in cells, due to the large burden of DNA required to be delivered into the cells.

The emergence of CRISPR/Cas technology has revolutionized the field of molecular biology, providing a promising tool for precise gene editing with profound implications for development of gene therapy ^1^. CRISPR/Cas utilises a RNA-guided mechanism for site-specific DNA cleavage, which has been used to knockin or knockout genes *in vitro* ^2,3^ and *in vivo* ^4,5^. However, the first described SpCas9 is large in size and presents a challenge for packaging in adeno associated viruses (AAV) for therapeutic delivery. Subsequently, smaller Cas9 variants have been identified, including SaCas9 ^6,7^, NmCas9 ^8^ and CjCas9 ^9^, which have higher therapeutic potential due to their smaller sizes. Also, further modifications of Cas9 have expand the ability of using CRISPR/Cas beyond genome editing, including control of gene regulation, epigenetics and chromatin imaging ^10^. For instances, a nuclease-null dead Cas9 can be fused to transcriptional activators to target the regulatory region of a gene to induce its expression ^11^. Importantly, multiple sgRNAs targeting different genes can be utilised to induce multiplex gene expression. Since only a short sgRNA is needed to induce expression of a single gene, rather than a whole transgene cDNA copy, the CRISPR activation approach can potentially reduce the number of viral vectors needed for overexpressing multiple genes. Another advantage of CRISPR activation is it activates gene expression at endogenous locus, which overcome the limitation of conventional gain-of-function studies that overexpress transgene beyond physiological levels and endogenous regulation.

Several reports have previously utilised a cloning-free CRISPR approach to knockin or knockout genes. In fission yeasts, Zhang *et al.* showed that the gap-repair mechanism can be used to assemble PCR-amplified sgRNA fragments and linear Cas9 plasmids together ^12^. In mammalian cells, Arbab *et al.* have reported a self-cloning CRISPR/Cas9 approach by using palindromic sgRNA, either in expression cassette or short DNA sequences, to allow homologous recombination in target cells to yield a functional site-specific sgRNA plasmid ^13^. The authors also showed that co-transfection of the Cas9 expression plasmids and sgRNA expression cassettes synthesized as a short 500bp oligonucleotides allow efficient knock out of target gene in mouse embryonic stem cells. Others have described the use of Cas9 protein complexes together with dual tracrRNA-crRNA for generation of knockin mice ^14,15^. However, a cloning-free approach has not been systematically reviewed for gene activation or repression using CRISPR/Cas9.

Here we present a method to use CRISPR/Cas for gene activation that is cloning-free. Our method utilises commercially synthesized sgRNA expression cassettes, which bypass the need for molecular cloning of site-specific sgRNA plasmids. In this study we compare the use of synthesized sgRNA expression cassessttes with two CRISPR/Cas activators, dSpCas9VPR and dSaCas9VPR, to induce gene expression in human and rat cells. Our strategy vastly simplifies the initiation of gain-of-function studies and as such has implications across many disciplines in cell biology.

## Results

To utilise CRISPR/Cas to activate endogenous genes in mammalian cells, we first tested the dSpCas9VPR system ^16^. This system utilises a nuclease-null dead SpCas9 (dSpCas9) coupled with transcription factor activation domains VP64, p65 and Rta (VPR). sgRNA were designed using the SAM sgRNA design tool. For gene activation, we chose six human genes that encoded for transcription factors that regulate neural and retinal development: *RAX, OTX2, RORB, CRX, ASCL1* and *NEUROD1* (Table 1). We synthesized sgRNA expression cassettes as linear oligonucleotide fragments containing an upstream U6 promoter, the sgRNA and the corresponding sgRNA scaffold (Supplementary figure 1). The sgRNA expression cassettes and the dSpCas9VPR plasmids were co-transfected into HEK293A cells. By day 3 post-transfection, we can detect robust gene activation in HEK293A cells using dSpCas9VPR, with robust upregulation of *OTX2* (∼375 fold increases, Figure 1A), *RAX* (∼13 fold increases, Figure 1B), *RORB* (∼110 fold increases, Figure 1C) and *CRX* (∼35 fold increases, Figure 1D). Similarly, the dSpCas9VPR system is also efficient in gene activation of *ASCL1* and *NEUROD1*, with ∼70 fold increases and ∼585 fold increases respectively (Figure 1E).

**Table 1:**
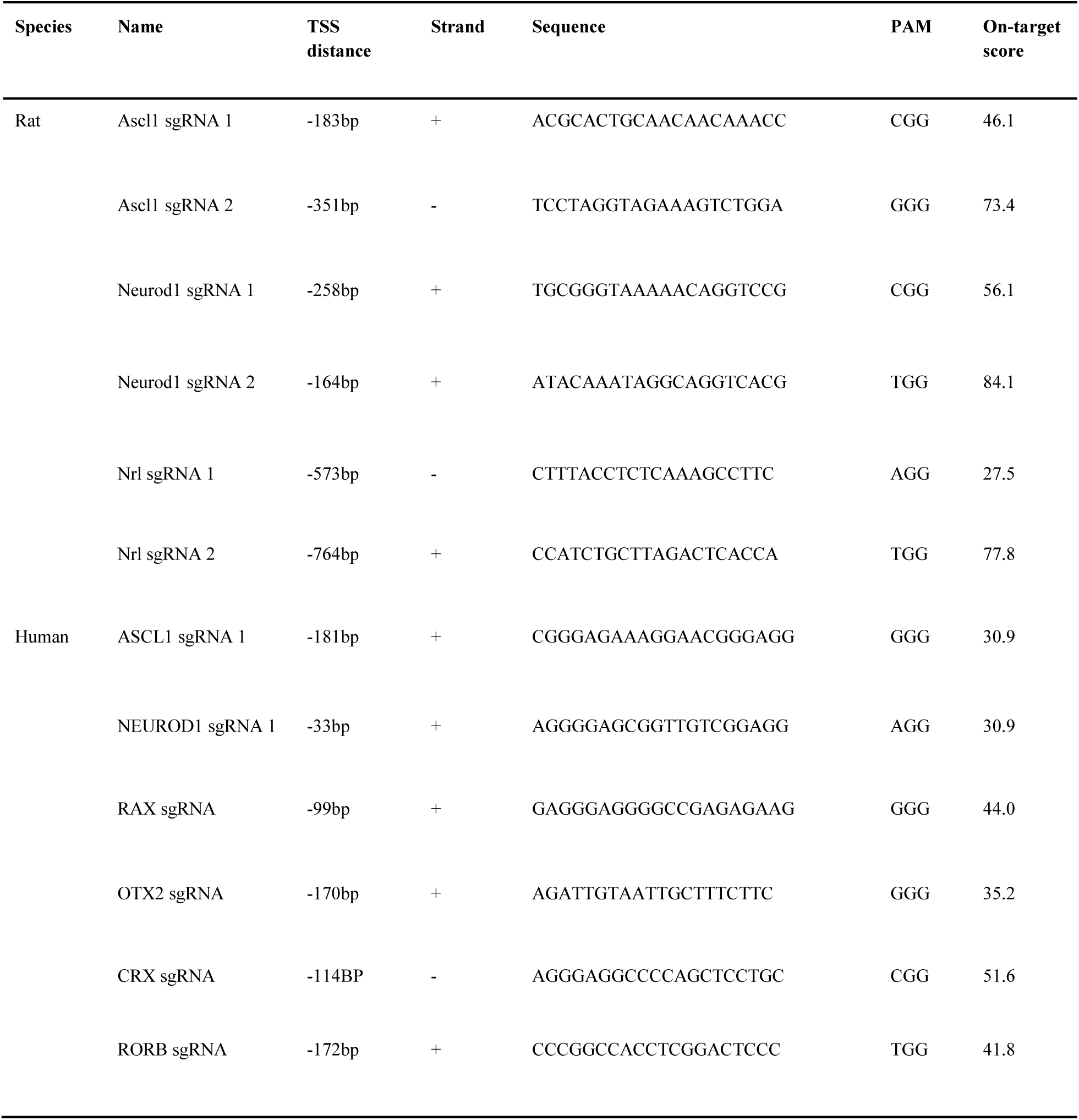
Information of sgRNAs used for SpCas9. On-target score is an optimized score for 20bp sgRNA based on ^17^. Scores range from 0-100. TSS: Transcription start site.

**Figure 1:**
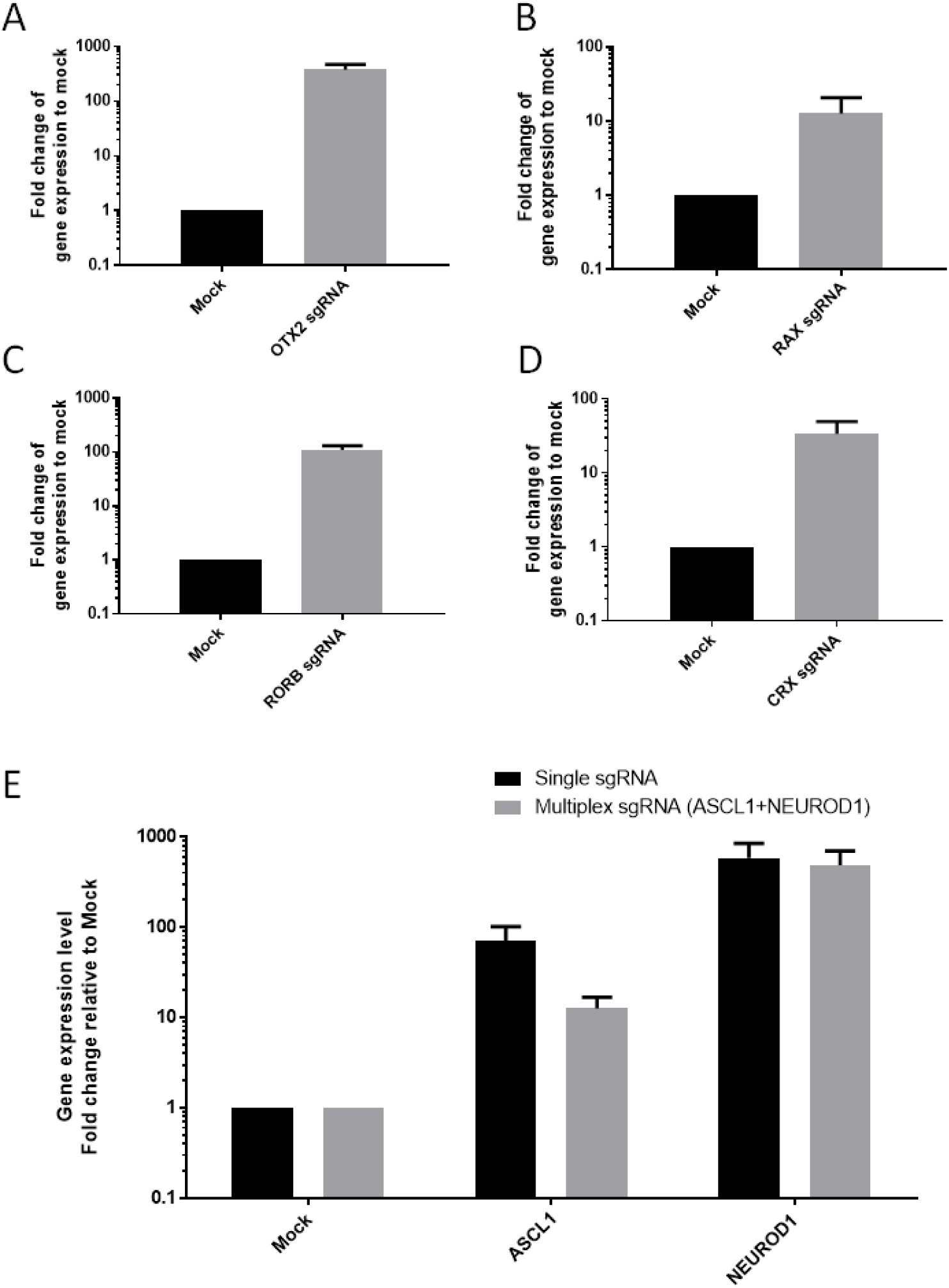
Efficient gene activation using dSpCas9VPR to induce gene activation in HEK293A cells. qPCR analysis of gene activation for A) *OTX2*, B) *RAX*, C) *RORB*, D) *CRX*, E) *ASCL1, NEUROD1* or multiplex induction of *ASCL1* and *NEUROD1*. Results are displayed as mean of three biological repeats ± SEM.

Subsequently, we tested the feasibility for using dSpCas9VPR for multiplex gene activation. We co-transfected sgRNAs for *ASCL1* and *NEUROD1* into HEK293A cells. Notably, both genes can be upregulated simultaneously and efficiently using the dSpCas9VPR, with ∼13 fold induction of *ASCL1* expression and ∼485 fold induction of *NEUROD1* expression (Figure 1E). However, multiplexing resulted in a lower level of gene induction compared to using single sgRNA, an effect more prominent in induction of *ASCL1* (Figure 1E). Taken together, our results demonstrated that dSpCas9VPR can be used to efficiently activate gene expression in human cells.

Furthermore, we assessed the feasibility of using dSpCas9VPR for gene activation in rat cells. As the SAM sgRNA design tool does not support design for the rat genome, we utilised the gene activator sgRNA design tool in Benchling (Table 1). We analysed dSpCas9VPR-mediated activation of three genes, *Ascl1, Neurod1* and *Nrl*, in a rat müller glial cell line rMC1, which can be transfected efficiently (Supplementary figure 4). Our results showed that *Ascl1* can be activated to modest levels using two sgRNAs, resulting in ∼3 and ∼2 fold increases respectively (Figure 2A). Similarly, two sgRNAs were tested for *Neurod1* gene activation. While one sgRNA resulted in a modest increase in *Neurod1* expression levels (∼3 fold increases), another sgRNA failed to activate *Neurod1* (Figure 2B). For *Nrl*, the two designed sgRNAs resulted in ∼3 and ∼1.6 fold induction in gene expression respectively (Figure 2C). We also tested gene activation with dSpCas9VPR in another rat fibroblast cell line R12. However, we did not observed significant gene activation in *Ascl1, Neurod1* nor *Nrl* (Supplementary figure 3). This is unlikely due to problems with transfection, as R12 can be efficiently transfected in this context (Supplementary figure 4). Collectively, these results demonstrated that dSpCas9VPR can be used to activate genes in rat cells, albeit only limited to modest levels of upregulation in certain rat cells and is not as efficient as in human cells.

**Figure 2:**
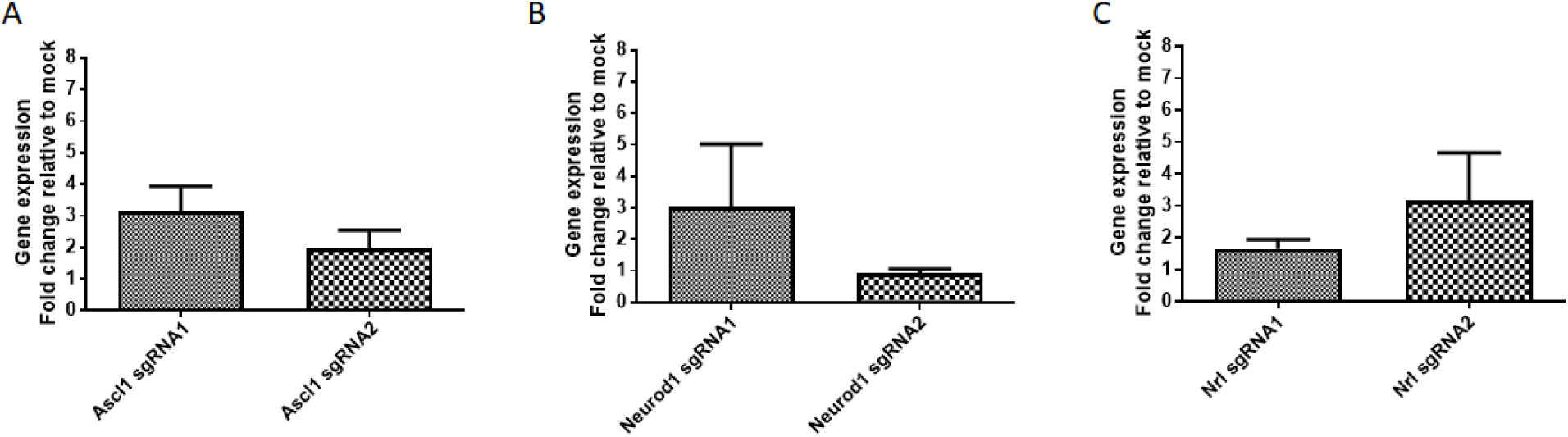
Assessment of multiple sgRNAs for dSpCas9VPR to induce gene activation in rat müller glial rMC1 cells. qPCR analysis of gene activation using 2 sgRNAs respectively for A) *Ascl1*, B) *Neurod1* and 3) *Nrl*. Results are displayed as mean of four biological repeats ± SEM.

Next, we assessed the efficiency of gene activation using another CRISPR/Cas activation system, dSaCas9VPR, which utilises a nuclease-null SaCas9 coupled with VPR. Notably, the dSaCas9VPR is much smaller in size than dSpCas9VPR, which have important implications for packaging into viral vectors for gene therapy. We designed sgRNAs using Benchling for both human and rat genes, and selected 2-3 sgRNAs for each gene for evaluation (Table 2). The sgRNA expression cassette for dSaCas9VPR contained a 5’ myc tag, upstream U6 promoter, sgRNA, sgRNA scaffold and a 3’ HA tag (Supplementary figure 2). In HEK293 cells, we showed that dSaCas9VPR can activate the expression levels of endogenous *ASCL1* efficiently (∼760 fold increases, Figure 3). However, dSaCas9VPR exhibited variable efficiency in gene activation in rMC1 cells (Figure 4), a result similar to those observed in dSpCas9VPR. Two sgRNAs were tested for activation of *Ascl1* or *Neurod1*, with 1 sgRNA inducing modest level of gene upregulation and the other fail to do so (*Ascl1:* ∼2.6, ∼1 fold changes; *Neurod1*: ∼2.8, ∼1 fold changes; Figure 4A, 4B). However, for *Nrl* activation all three sgRNA tested failed to upregulate its expression levels (Figure 4C). In summary, we showed that dSaCas9VPR can be used to efficiently activate gene expression in human cells, however we found its effects in rat cells is variable and remained inefficient.

**Table 2:**
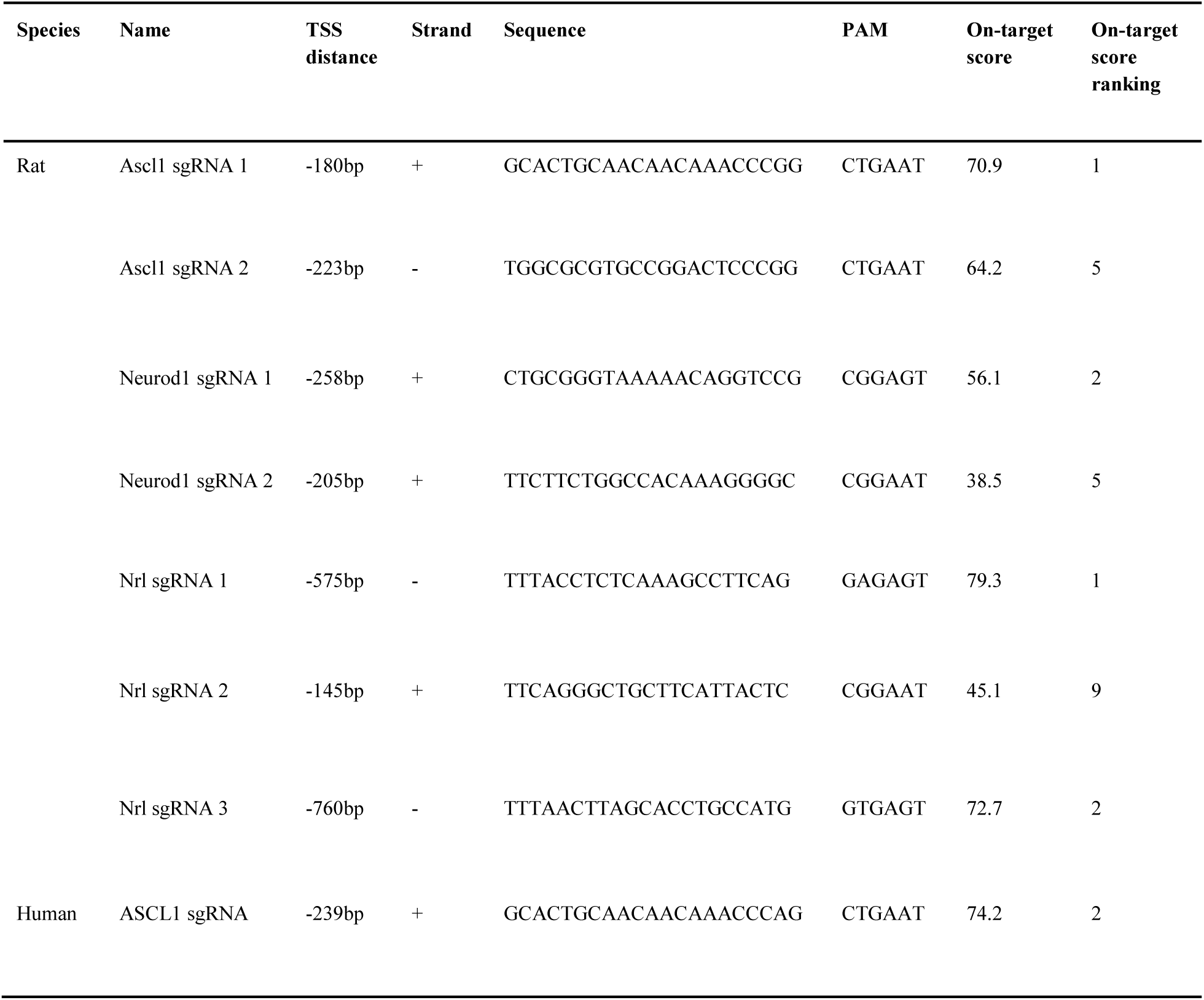
Information of sgRNAs used for SaCas9. On-target score is an optimized score for 20bp sgRNA based on ^17^. The on-target scores are ranked for sgRNAs targeting 2000bp upstream of transcription start site of genes. Scores range from 0-100. TSS: Transcription start site.

**Figure 3:**
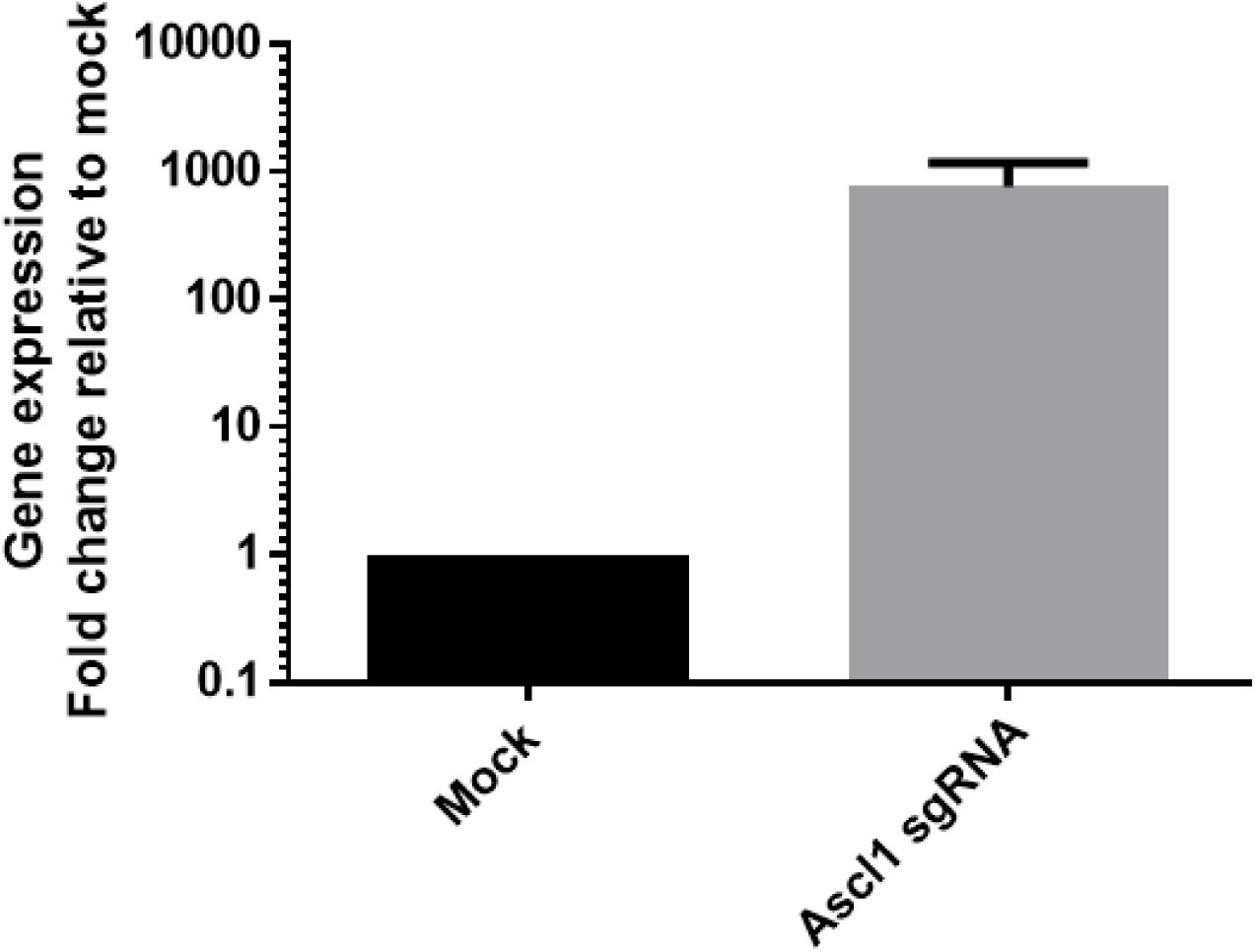
Efficient gene activation using dSaCas9VPR in HEK293A cells. qPCR analysis of gene activation for *ASCL1* in HEK293 cells. Results are displayed as mean of three biological repeats ± SEM.

**Figure 4:**
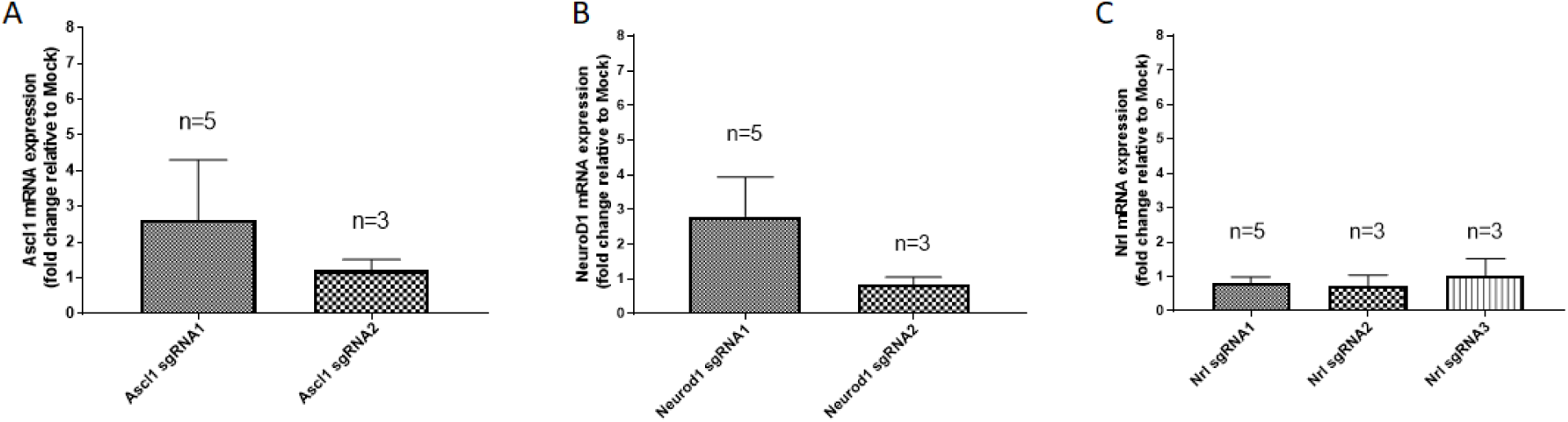
Comparison for multiple sgRNAs for dSaCas9VPR to induce gene activation in rat müller glial rMC1 cells. qPCR analysis of gene activation for A) *Ascl1* using 2 sgRNAs, B) *Neurod1* using 2 sgRNAs and 3) *Nrl* using 3 sgRNAs. Results are displayed as mean of 3-5 biological repeats ± SEM.

## Discussion

A key limiting step in the conventional design of overexpression studies is the requirement of tedious plasmid construction for each transgene, which involves molecular cloning steps that take >1 week. Here we present a simplified method for gene activation using CRISPR/Cas9 activators in mammalian cells that is feasible in one day. Notably, our strategy to transfect sgRNA as a synthesized linear oligonucleotide fragment with U6 promoter allow sgRNA expression in the cells, and eliminates the need to clone sgRNA into a designated vector prior to transfection. By cutting down the time required for molecular cloning, our strategy provides a rapid way to initiate gene overexpression studies using the CRISPR activation system.

This study compared two CRISPR activation systems, dSpCas9VPR and dSaCas9VPR, in human and rat cells. Our results demonstrated that both dSpCas9VPR and dSaCas9VPR can efficiently induce gene expression in human cells. We also showed the use of dSpCas9VPR for multiplex activation of two genes. In rat cells, in total we have assessed activation of three genes, using six sgRNAs for dSpCas9VPR and seven sgRNAs for dSaCas9VPR, in two different rat cell lines. However, our results showed only modest levels, and in some cases, negligible levels of gene activation in rat müller glial cells and fibroblasts. This is unlikely due to issues with transfection efficiency as both rat cell types can be efficiently transfected with robust GFP expression. We speculate that this is due to inadequate support of current sgRNA design algorithm for the rat genome. For instances, many of the sgRNAs used for rat genes have high ranking of on-target scores (Table 2), but yet they are mostly inefficient in inducing expression of rat genes using the CRISPR activation systems. This potentially highlights the need to better design and improve accuracy of predicting functional sgRNAs for rat genome.

A potential challenge of using CRISPR/Cas systems is that not all sgRNAs are efficient in Cas9 targeting, thus multiple sgRNAs are often screened to identify the functional sgRNAs. In our experience, the sgRNA design tools for predicting functional sgRNA are generally very accurate for human cells. For dSpCas9VPR, the first sgRNA designed is functional for gene activation in 13 out of 15 genes tested (∼87%, Supplementary table 1). Similarly, although we only tested gene activation for a single gene using dSaCas9VPR, the first sgRNA designed is highly efficient for gene activation (Table 3). Therefore, we recommend designing two sgRNAs within 300bp upstream of transcription start site to be tested for gene activation in human cells.

In summary, this study outlines a simple and robust workflow to efficiently activate endogenous gene expression in mammalian cells using CRISPR/Cas activators, which can be applied as a rapid workflow to initiate gene overexpression studies for a range of molecular and cell biology subjects.

## Materials and methods

### sgRNA design and preparation of expression cassette

For SpCas9, sgRNAs with NGG PAMs were designed using the SAM sgRNA design tool (http://sam.genome-engineering.org/database/) for human genes, and designed using Benchling (https://benchling.com/) for rat genes. The SpCas9 sgRNA expression cassette contains an upstream U6 promoter, sgRNA and sgRNA scaffold with stem extension and stem loop (Supplementary figure 1).

For SaCas9, sgRNAs with NNGRRT PAMs were designed using Benchling (https://benchling.com/). Predicted sgRNAs with long stretches of repeating nucleotides are excluded from selection. The SaCas9 sgRNA expression cassette contains a Myc tag, U6 promoter, sgRNA, sgRNA scaffold and HA tag (Supplementary figure 2).

Both SpCas9 and SaCas9 sgRNA expression cassettes (<500bp) were synthesized as gBLOCK gene fragments (Integrated DNA Technologies). The sgRNA expression cassettes were amplified by PCR using the following primers: SpCas9 forward primer: TGAGTATTACGGCATGTGAGGGC,SpCas9reverseprimer: TCAATGTATCTTATCATGTCTGCTCGA;SaCas9forwardprimer: GAACAAAAACTCATCTCAGAAGAGGATCTG, SaCas9 reverse primer: TACCCATACGATGTTCCAGATTACGCT. PCR were performed using KOD Hot Start DNA polymerase (Merck Millipore), with the following thermal profile: 95°C for 2 minutes; 30 cycles of 95°C for 20 seconds, 66°C (SpCas9 sgRNA) or 64°C (SaCas9 sgRNA) for 10 seconds, 70°C for 8 seconds; 70°C for 5 minutes. The PCR amplicons were separated by gel electrophoresis and the sgRNA expression cassettes were extracted using the Wizard SV gel and PCR cleanup kit (Promega). The amplified sgRNA expression cassettes were checked with Nanodrop to confirm good DNA quality.

### Cell culture

rMC1 rat muller glia, R12 rat fibroblasts and HEK293A cells were maintained in DMEM high glucose media supplemented with 10% fetal calf serum, 2mM L-glutamine, 0.5% penicillin/streptomycin (all from ThermoFisher). All cells were passaged using 0.25% trypsin before the culture become confluent, and maintained in incubators at 37°C with 5% CO2 level.

### Transfection efficiency assay

rMC1 and R12 cells were transfected with the pmaxGFP construct (Lonza) using Lipofectamine 3000 overnight, following the manufacturer’s instruction. GFP expression is determined 1 day after transfection using a fluorescence microscope (Olympus CKX53).

### Gene activation using CRISPR/Cas activation

dSaCas9-VPR and dSpCas9-VPR plasmids were gifts from George Church (Addgene #68495 and #63798 respectively). rMC1, R12 and HEK293A cells were transfected using Lipofectamine 3000. Briefly, cells with were plated down on a 12 well plate at day 0 (rMC1, HEK293: 6 x 10^4^/well). At Day 1, the cells were transfected with 360ng sgRNA expression cassettes and 800ng dSaCas9-VPR or dSpCas9-VPR plasmids using Lipofectamine 3000 overnight. In some experiments the cells and DNA were upscale proportionally to obtain more RNA. At day 4, the samples were harvested for RNA to assess gene expression levels.

### qPCR analysis

Total RNA was extracted using the RNeasy kit (Qiagen) or the Illustra RNAspin kit (GE Healthcare) followed by DNase treatment. cDNA synthesis was performed using the High capacity cDNA Reverse Transcription Kit (ThermoFisher) following the manufacturer’s instructions. Taqman gene expression assay (ThermoFisher) was performed using the Taqman Fast Advanced Master Mix with the following probes: human *RAX* (Hs00429459_m1), human *OTX2* (Hs00222238_m1), human *ASCL1* (Hs00269932_m1), human *NEUROD1* (Hs00159598_m1), human *CRX* (Hs00230899_m1), human *RORB* (Hs00199445_m1), human *ACTB* (Hs99999903_m1), rat *Ascl1* (Rn00574345_m1), rat *Neurod1* (Rn00824571_s1), rat *Nrl* (Rn01481925_m1) and rat *Gapdh* (Rn01775763_g1). qPCR was processed using the ABI StepOnePlus system or the ABI 7500 system (Applied Biosystems). The delta delta Ct method was used to calculate relative gene expression to control. The housekeeping genes *ACTB* and *Gapdh* were used for normalising gene expression in human and rat cells respectively.

## Acknowledgement

We thank Chris Karelas for technical support. rMC1 is a generous gift from Mark Gillies’s laboratory and R12 is a generous gift from Peter Van Wijngaarden. This work was supported by funding from the Ophthalmic Research Institute of Australia (RW), the University of Melbourne De Brettville Trust (RW) and the Kel and Rosie Day Foundation (RW). The Centre for Eye Research Australia receives operational infrastructure support from the Victorian Government.

## Authors contributions

SH, AWH and RW designed the experiments; LF, JY, TN, SK, SH performed the experiments; LF, JY, SH, AWH, RW analysed the data; RW wrote the manuscript; all authors approved the manuscripts.

## Supplementary information

**Supplementary figure 1:**
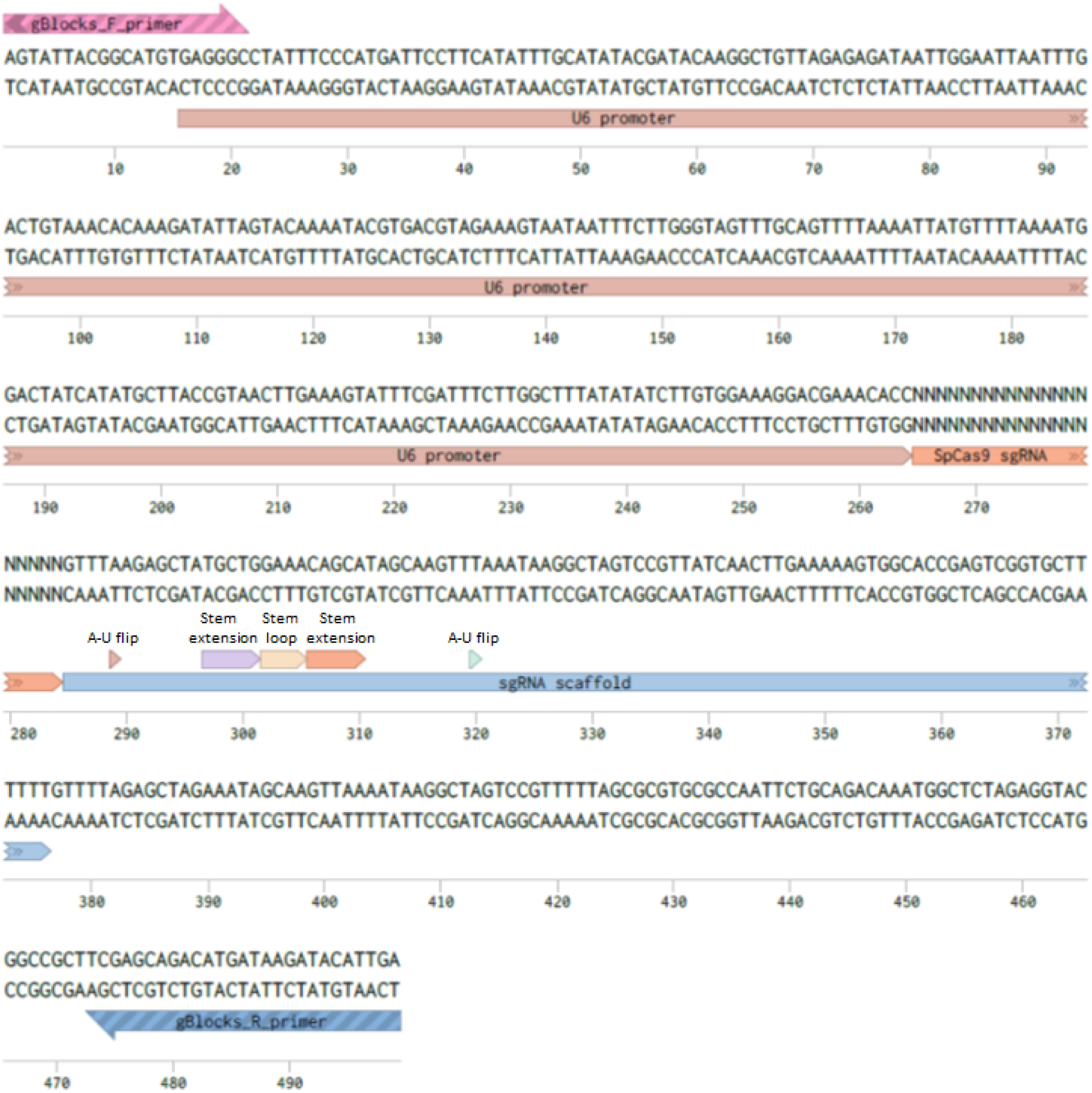
Design of sgRNA expression cassette for SpCas9.

**Supplementary figure 2:**
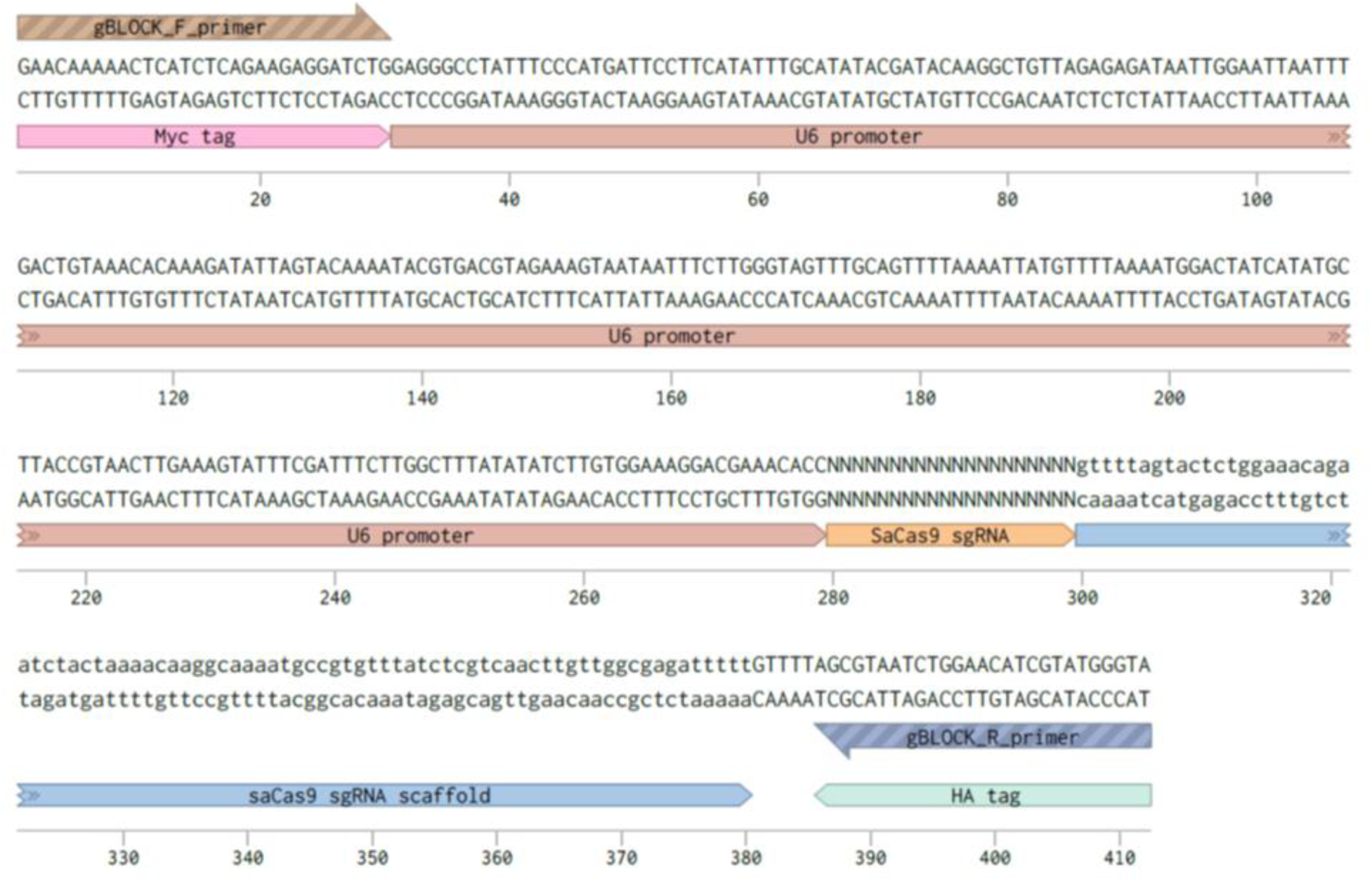
Design of sgRNA expression cassette for SaCas9.

**Supplementary figure 3:**
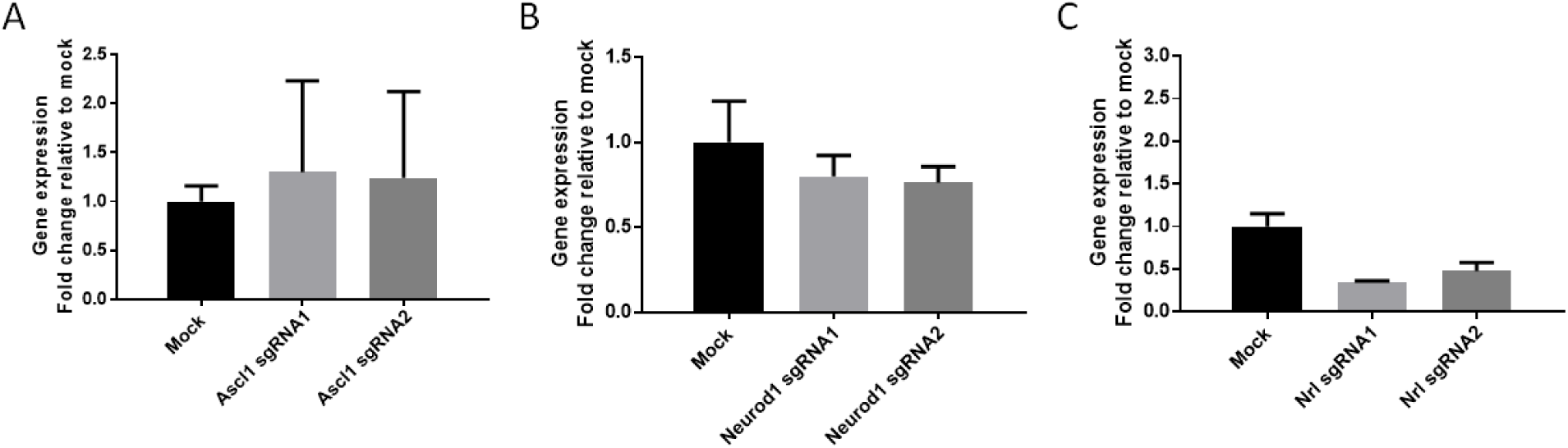
Assessment of multiple sgRNAs for dSpCas9VPR to induce gene activation in rat fibroblast R12 cells. Results are displayed as mean of three technical repeats ± SEM.

**Supplementary figure 4:**
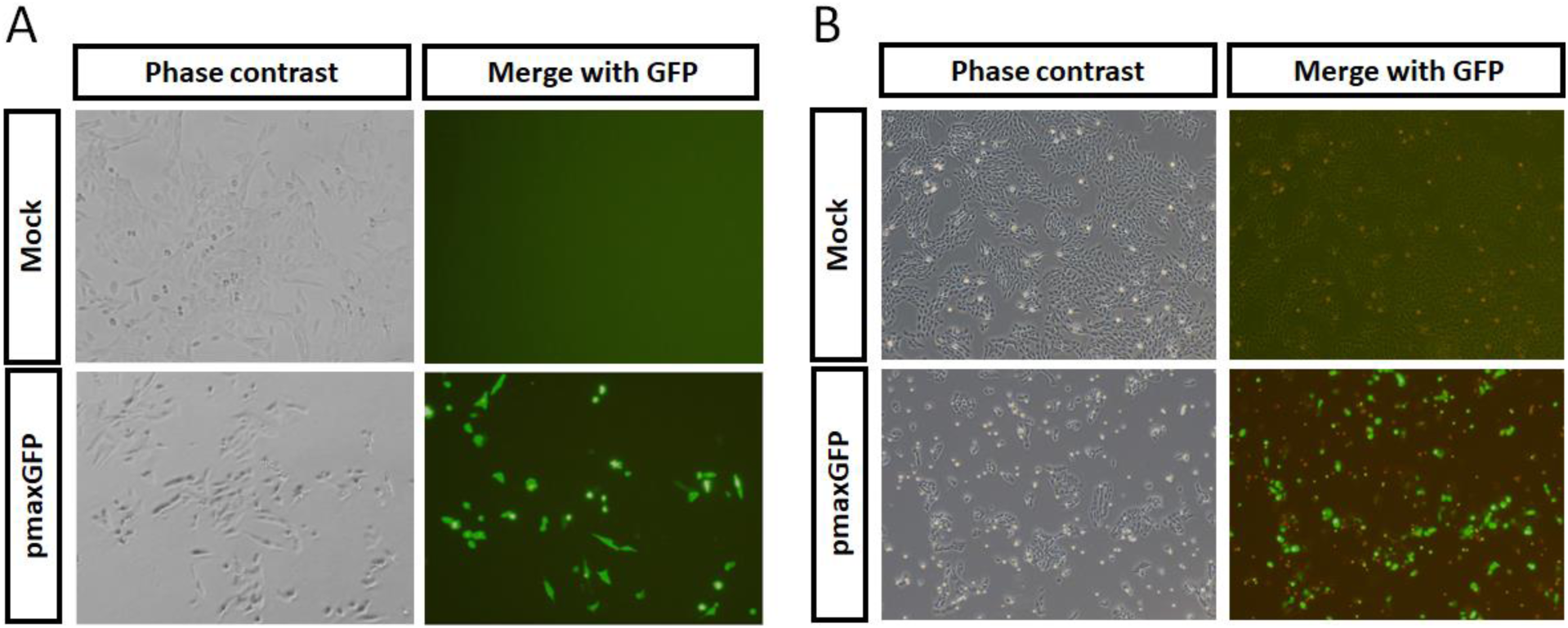
Lipofectamine 3000 allows efficient transfection in A) rMC1 rat müller glial and B) R12 rat fibroblasts.

**Supplementary table 1:** Success rate of designing sgRNAs for SpCas9 and SaCas9 in human cells

| **Cas9 activator** | **Species** | **Genes/sgRNA** | **Number of genes** | **Success rate** |
| --- | --- | --- | --- | --- |
| dSpCas9VPR | Human | Total genes tested | 15 | 100% |
|  |  | Gene activation with first sgRNA | 13 | 86.7% |
|  |  | Gene activation with second sgRNA | 1 | 6.7% |
|  |  | Gene activation with third sgRNA | 1 | 6.7% |
| dSaCas9VPR | Human | Total genes tested | 1 | 100% |
|  |  | Gene activation with first sgRNA | 1 | 100% |

